# Bacteriophage Heteroresistance as a Cause of Treatment Failure in Urinary Tract Infections

**DOI:** 10.64898/2026.06.26.734749

**Authors:** Serena Singh-Ward, Alysha S. Ismail, Teresa Gil-Gil, Brandon A. Berryhill, Michael H. Woodworth, Hope E. Shanks, Bruce R. Levin

## Abstract

With the rise of antimicrobial resistance, urinary tract infections (UTIs) have become increasingly more difficult to treat, prompting renewed interest in bacteriophage (phage) therapy as an alternative or adjunct to antibiotics. UTIs are an attractive target for phage therapy because they generate a high density of actively replicating bacteria that supports phage propagation, and because the urinary tract is readily accessible for administration and monitoring. Yet studies of phage therapy for UTIs report mixed outcomes, including failures to meet clinical and microbiological endpoints.

Here we follow the population dynamics of a clinical *Escherichia coli* UTI strain and two phages, HP3 and ES19, to which the strain appears susceptible by standard testing. Despite this apparent susceptibilty, both phages fail to suppress the strain, with resistance emerging almost immediately. Using the measured mutation rate, our mathematical model shows that traditional resistance cannot account for these dynamics. We instead demonstrate, including by a phage-specific population analysis profile assay we developed, that heteroresistance drives this rapid failure, offering a plausible explanation for treatment failures in UTI phage therapy.

## Introduction

Urinary tract infections (UTIs) are among the most common bacterial infections with approximately 10% of women in the USA experiencing a UTI each year, and nearly half of those patients experiencing a recurring UTI within the same year^1^. Historically, these infections have been straightforward to diagnose and treat, but rising antibiotic resistance has complicated UTI management^2,3^, renewing interest in bacteriophage (phage) therapy^4^. UTIs are an especially attractive candidate for phage therapy for several reasons: (1) These bacterial infections typically produce a high density of actively growing bacteria, creating the ideal environment for phage to flourish^5,6^; (2) There is ease of access to the infection site for diagnosis and administration of phage therapy^7^; and, (3) UTIs are commonly caused by uropathogenic *Escherichia coli* (UPEC) which are one of the most well studied bacterial pathogens^8^.

Numerous studies of phage therapy for UTIs report mixed treatment outcomes. Some of these studies have shown favorable results, including complete clearance of the infection ^9^ or the patient becoming asymptomatic^10^. Conversely, other case studies resulted in unfavorable outcomes where either the infection became recurrent or *in vitro* testing of the phage did not translate into *in vivo* success^9^. The conventional explanation for phage therapy failure is the evolution of resistance.

Here, we study a representative UPEC strain, *E. coli* DS566, and two phages, HP3 and ES19, that have been used in UTI phage therapy^10^. *E. coli* DS566 was isolated from a patient with neurogenic bladder due to spinal cord injury^11^. The phages HP3 and ES19 were isolated from duck waste samples and wastewater, respectively^12^. These phages are not fully characterized; therefore, their specific bacterial receptors remain uncertain^11^. Sequencing data from a previous study shows mutations occurring in either the lipopolysaccharides (LPS) or OmpA protein of an *E. coli* strain when exposed to HP3, suggesting that the receptor for HP3 may be the LPS and/or OmpA^13^; whereas ES19’s receptor is unverified but is thought to be an LPS-associated moiety^14^. Using susceptibility testing methods, both phages appear capable of controlling the growth of *E. coli* DS566^15,16^. However, by assessing the population and evolutionary dynamics of these phages, we observe that neither fully suppresses the growth of this bacterium. This failure is not due to standard resistance, but rather phage heteroresistance. The overarching indicator of heteroresistance is that these bacterial strains appear susceptible by the spot test assay, but upon exposure to the phage, minor resistant populations rapidly ascend to dominance. Importantly, we found that this resistance can be unstable, with some lineages reverting to susceptibility^17^. Overall, heteroresistance is largely under described in the field of phage therapy and may account for treatment failures in UTI phage therapy^18,19^.

## Results

### HP3 and ES19 are suitable candidates for treating UTIs

To evaluate the effectiveness of phage therapy on the treatment of UTIs, we challenged the clinical UTI strain *E. coli* DS566 with two distinct phages, HP3 and ES19, in pooled human urine media. The Optical Density (OD) suppression time of these phages was measured to be 8.46 ± 1.56 and 10.13 ± 1.27 hours at an MOI of 10, respectively. A spot test of *E. coli* DS566 indicates that this strain is susceptible to both phages (Supplemental Figure 1).

### Phage resistance rapidly emerges

To observe the dynamics of *E. coli* DS566 in the presence of HP3 and ES19 in urine, we performed a time kill assay (Figure 1). Within an hour, *E. coli* DS566 rapidly declines before resistant cells increase in frequency, and, by two hours, resistant cells take over in both HP3- and ES19-treated cultures. By 24 hours, the *E. coli* DS566 cells that were exposed to phage reach the same final density as the control group. Interestingly, the bacterial population only declines to about 1/100^th^ of the starting bacterial concentration before the resistant subpopulation ascends, indicating the presence of a large phage-resistant subpopulation. Qualitatively similar results were obtained in LB (Supplemental Figure 2).

**Fig 1.**
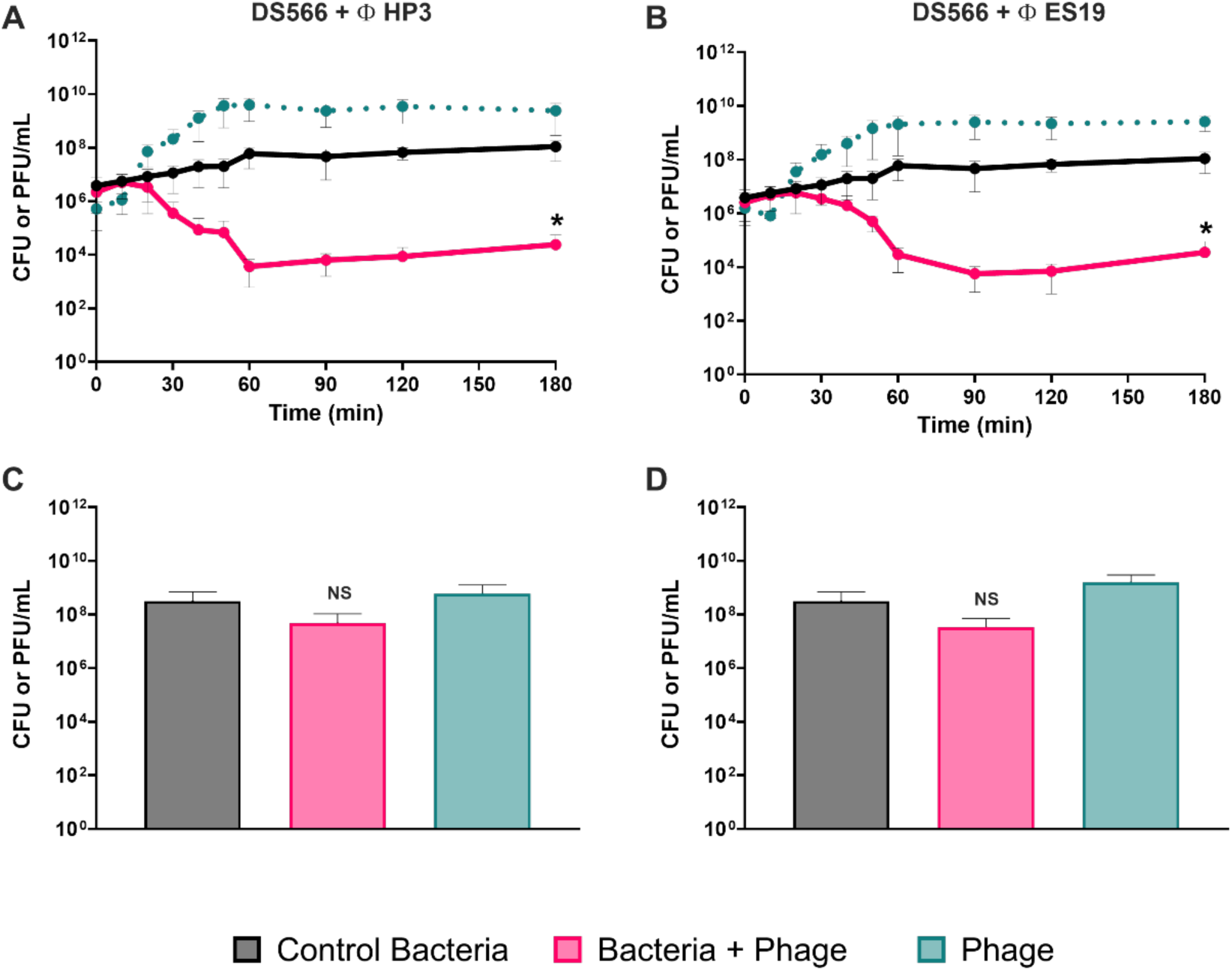
Time kills of *E. coli* DS566 with either phage in urine. Shown at the top are three-hour time kills with *E. coli* DS566 and HP3 **(A)** or ES19 **(B)**. Shown below are the 24-hour densities from the time kill experiments **(C, D)**. In black is the bacterial density of a phage free control, pink is the bacterial density of a culture with the phage, and blue is the phage density in the culture with the phage. Shown are means and standard deviations of three biological replicates. \**p* < 0.1, ns = not significant.

### Analysis of the resistant isolates

Resistant colonies were isolated between the one- and two-hour timepoints from the time kill experiments once the initial phage-sensitive population had been killed. Based on the spot test, these mutants were confirmed to be resistant to both phages—even after exposure to just one phage in the time kill (Supplemental Table 1). This cross-resistance is likely due to mutations in the LPS as both phages likely target different moieties on LPS^11^ (Supplemental Table 2).

Growth curve data of all 16 phage-resistant colonies collected revealed variable lag times and fitness costs (Supplemental Figure 3). Out of all mutants screened, *E. coli* DS566-11 and *E. coli* DS566-15 were selected for further analysis as *E. coli* DS566-11 had almost no fitness cost for resistance and *E. coli* DS566-15 had the highest fitness cost in urine (Figure 2) and LB (Supplemental Figure 4).

**Fig 2.**
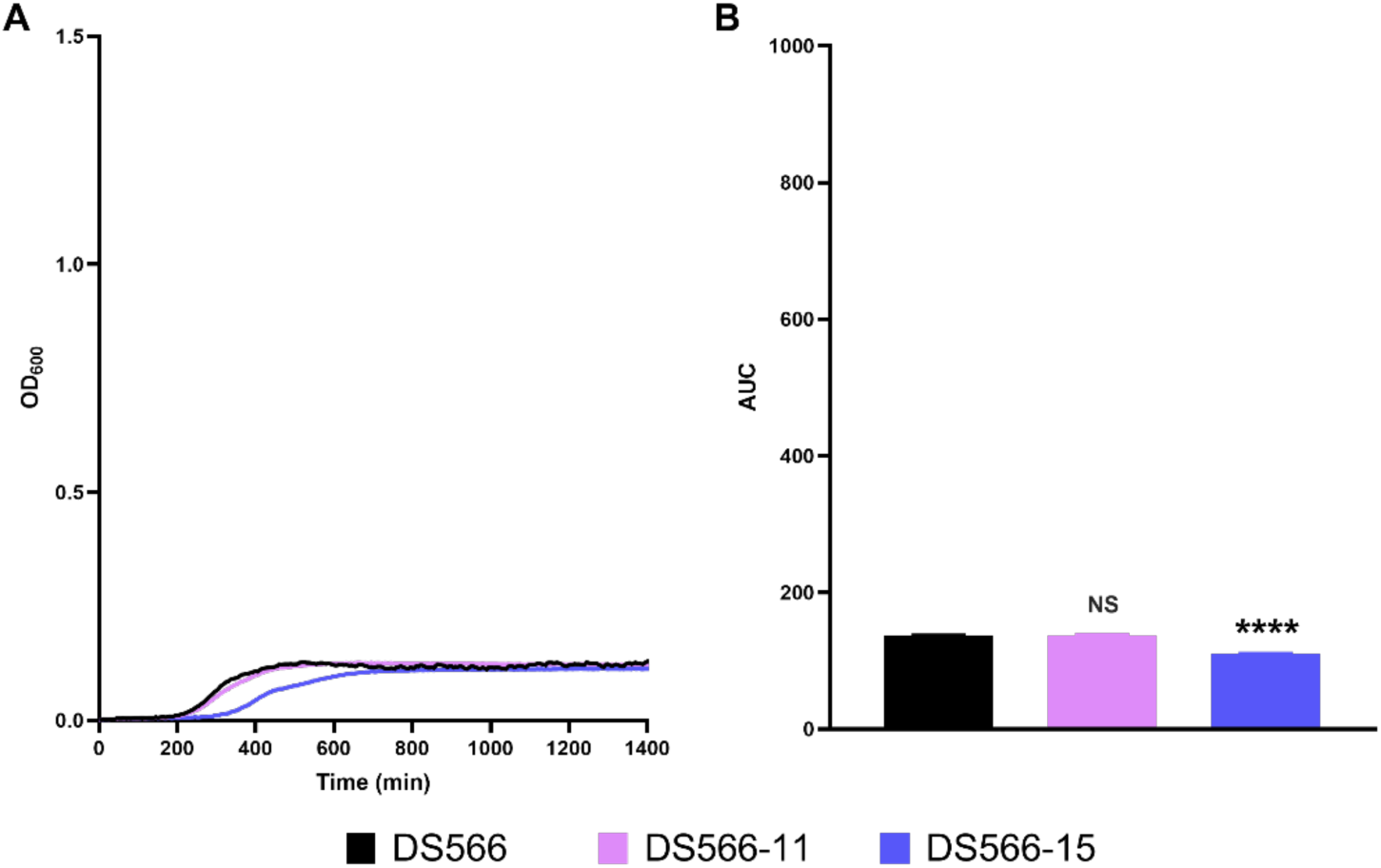
Fitness cost of phage-resistant mutants in urine. Shown in black is the ancestral *E. coli* DS566, in pink a low-fitness-cost mutant, and in indigo a high-fitness-cost mutant. **(A)** OD growth curves. **(B)** Area under the curve of the OD growth curves. \*\*\*\**p* < 0.0001, ns = not significant.

To observe the dynamics of these phage-resistant mutants in the presence of HP3 and ES19, we performed a time kill assay in urine (Figure 3) and LB (Supplemental Figure 5). Contrary to the time kills shown in Figure 1 and Supplemental Figure 2, *E. coli* DS566-11 and *E. coli* DS566-15 were completely resistant throughout the time kill experiment.

**Figure 3.**
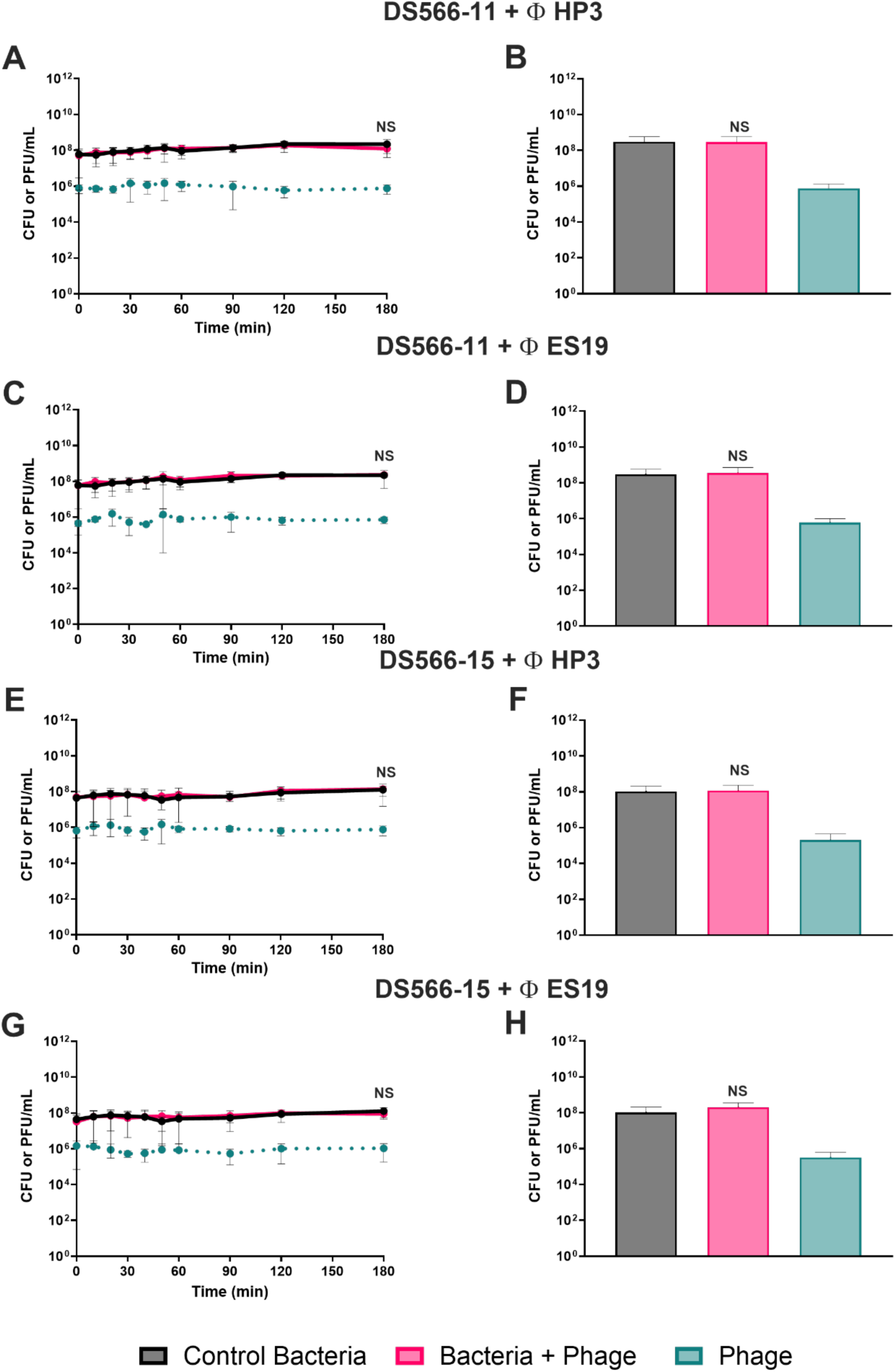
Time kills of phage-resistant mutants with either phage in urine. **(A)** Time kill with *E. coli* DS566-11 with the phage HP3. **(B)** 24-hour densities of *E. coli* DS566-11 with the phage HP3. **(C)** Time kill with *E. coli* DS566-11 with the phage ES19. **(D)** 24-hour densities of *E. coli* DS566-11 with the phage ES19. **(E)** Time kill with *E. coli* DS566-15 with the phage HP3. **(F)** 24-hour densities of *E. coli* DS566-15 with the phage HP3. **(G)** Time kill with *E. coli* DS566-15 with the phage ES19. **(H)** 24-hour densities of *E. coli* DS566-15 with the phage ES19. ns = not significant. In black is the bacterial density of a phage-free control, pink is the bacterial density of a culture with the phage, and blue is the phage density in the culture with the phage. Shown are means and standard deviations of three biological replicates.

### Standard mutational resistance is insufficient to explain the dynamics

To determine the mutation rate of *E. coli* DS566 to HP3 and ES19, we performed a Luria-Delbrück fluctuation test (Table 1). To ascertain whether the rapid rise of resistance was bacterial strain dependent and/or phage dependent, we first performed the same test against the antibiotic streptomycin and second, a fluctuation test with the same phages but against an unrelated strain of *E. coli*, *E. coli* A16.

**Table 1.** Mutation Rates.

| Phage or Antibiotic Treatment | DS566 Mutation Rate | A16 Mutation Rate |
| --- | --- | --- |
| **HP3** | $9.44 \times 10^{-7} \pm 3.3 \times 10^{-7}$ | $6.57 \times 10^{-7} \pm 4.27 \times 10^{-7}$ |
| **ES19** | $3.28 \times 10^{-7} \pm 1.17 \times 10^{-7}$ | $6.57 \times 10^{-7} \pm 3.75 \times 10^{-7}$ |
| **Streptomycin** | $4.51 \times 10^{-8} \pm 1.74 \times 10^{-8}$ | $2.51 \times 10^{-9} \pm 5 \times 10^{-1}$ |

Statistical analysis presented in Supplemental Table 3.

Exposure of *E. coli* DS566 to streptomycin yielded an approximately ten-fold lower mutation rate than to either phage. Additionally, *E. coli* A16 was found to have mutation rates similar to those of *E. coli* DS566 to both phages. These results indicate that the high mutation rate is determined by a property of the phage rather than by the bacterial strain (e.g., being a mutator strain). DS566 colonies taken from the HP3 fluctuation test were confirmed by spot test to be resistant to both phages.

To ascertain the effect that the rate of resistance generation has on the final bacterial density, we employed a mathematical and computer-simulation model of envelope resistance with the mutation rates obtained in Table 1 (Supplemental Text, Supplemental Equations 1–4, Supplemental Table 4). In Figure 4, we show that resistance cannot account for the population levels obtained in Figure 1 after three hours, such that in the time kill experiment, the total number of surviving cells would be lower if envelope resistance was the sole driver of the bacterial dynamics.

**Fig 4.**
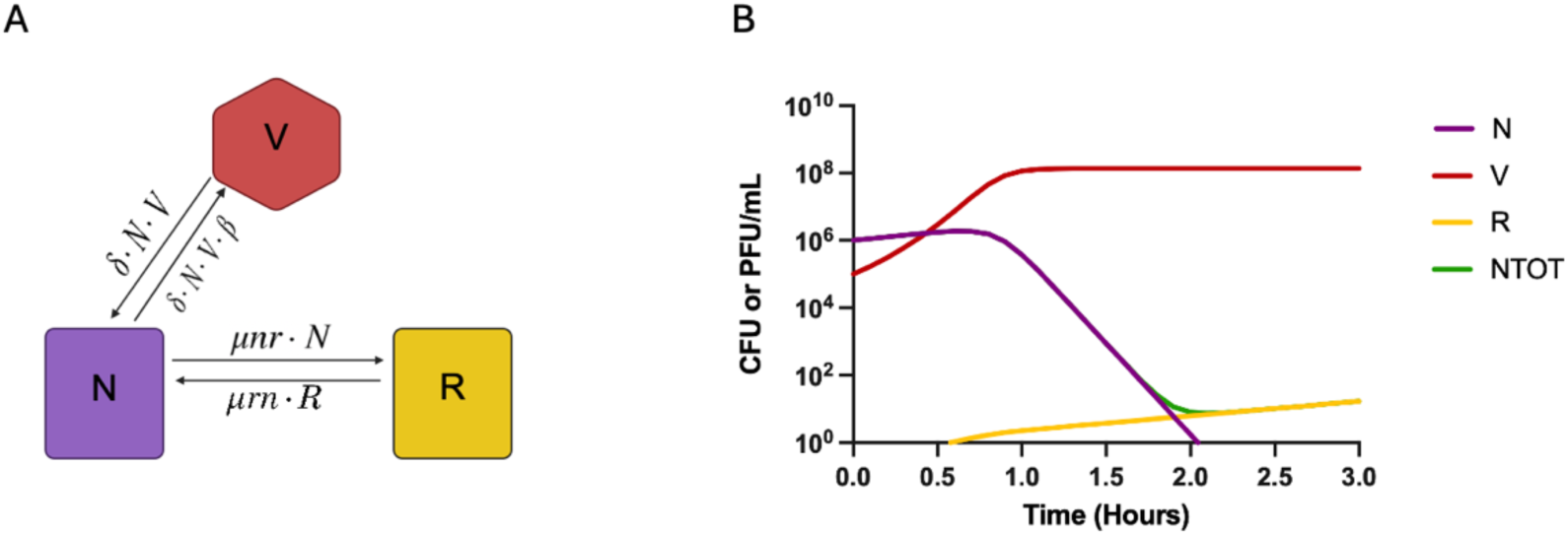
Diagram and Predictions of the model of envelope resistance. **(A)** Diagram of the envelope resistance model. The variables N, V, and R are, respectively, sensitive bacteria, viral particles, and resistant bacteria in cells or phage per mL. **(B)** Computer simulations of the proposed model of envelope resistance. N (sensitive bacteria, purple), V (viral particles, red), R (resistant bacteria, yellow), NTOT (total bacteria, green).

### Heteroresistance can explain the dynamics

Heteroresistance is defined as the occurrence of a subpopulation that is resistant within a larger susceptible population^20^. Traditionally, a population analysis profile (PAP) test is the standard assay for antibiotic heteroresistance. Currently, there is no method for testing phage heteroresistance as the PAP test is unsuitable.

The PAP test designed for antibiotics quantifies the resistant subpopulations over a range of antibiotic concentrations. This is done by plotting the number of surviving colonies at each antibiotic concentration to generate a curve. The strain is determined to be heteroresistant if there are subpopulations present at a frequency of greater than 10^−7^ with an MIC 8 times higher than the main population^21^. This method is only suitable for antibiotics because the number of surviving colonies is directly dependent on the antibiotic’s fixed concentration.

Unlike antibiotics, phages are self-replicating, which means the concentration of phage will increase over time from the initial concentration. Moreover, the time at which each bacterium is infected will vary. Hence, a surviving colony count cannot directly correlate with the initial phage MOI. Therefore, the PAP test must be redesigned to assess phage heteroresistance.

Similar to how the traditional PAP test can show the decline of susceptible cells and the survival of a resistant subpopulations at high antibiotic concentrations, our PAP test must be able to demonstrate this for phage. This is accomplished by inoculating wells with different initial densities of bacteria and a fixed density of phage. In the case of traditional resistance, we expect all wells to grow due to every cell in the population being resistant to the phage. For a bacterial population that is heteroresistant, we expect to see resistance only in wells which have cells from the resistant subpopulation (namely, those wells at or above the frequency of the resistant subpopulation). Since heteroresistance is the generation of resistant subpopulations at a high frequency, our PAP test is a test for the approximate size of the resistant subpopulation, which we expect to vary as these mutants emerge stochastically.

We performed our phage-specific test for heteroresistance against the ancestral *E. coli* DS566 and the phage-resistant mutants and determined the frequencies of the phage-resistant subpopulations (Figure 5 and Supplemental Figure 6). The PAP test for *E. coli* DS566 shows that an initial stationary-phase culture of this bacterium contains between 100 and 100,000 phage-resistant cells (Figure 5C and 5D and Supplemental Figure 6C and 6D). Notably, when the resistant clones selected from the time kills themselves are exposed to the phage, the entire population is resistant (Figure 5E-H and Supplemental Figure 6E-H). We used *E. coli* C and the phage T6 (Figure 5A and Supplemental Figure 6A), as well as *E. coli* DS566 with the phage T3 (Figure 5B and Supplemental Figure 6B) as susceptible and resistant controls, respectively. The susceptible control shows no growth, and the resistance control shows growth in all 96 wells of the plate.

**Fig 5.**
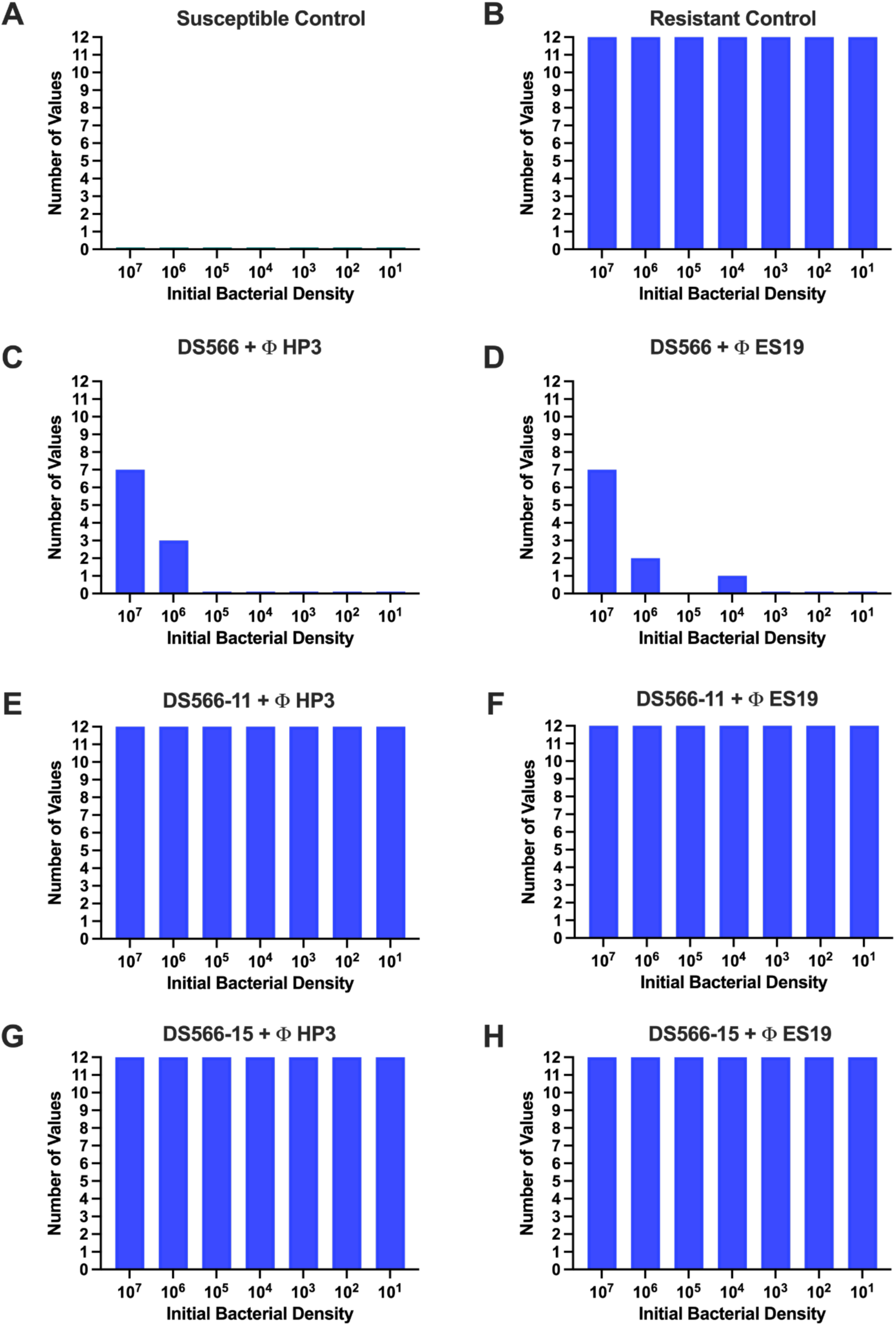
Phage PAP test in urine. **(A)** *E. coli* C with the phage T6 (susceptible control). **(B)** *E. coli* DS566 with the phage T3 (resistant control). **(C)** *E. coli* DS566 with the phage HP3. **(D)** *E. coli* DS566 with the phage ES19. **(E)** *E. coli* DS566-11 with the phage HP3. **(F)** *E. coli* DS566-11 with the phage ES19. **(G)** *E. coli* DS566-15 with the phage HP3. **(H)** *E. coli* DS566-15 with the phage ES19.

Next, to examine if heteroresistance can in fact account for the dynamics observed in Figure 1 and Supplemental Figure 2, we employed a mathematical and computer-simulation model of heteroresistance (Supplemental Text, Supplemental Equations 5–8, Supplemental Table 5). In Figure 6, we demonstrate that the predictions obtained with the heteroresistance models mirror the time kill results, providing evidence that heteroresistance can account for the observed dynamics in the presence of the phage.

**Fig 6.**
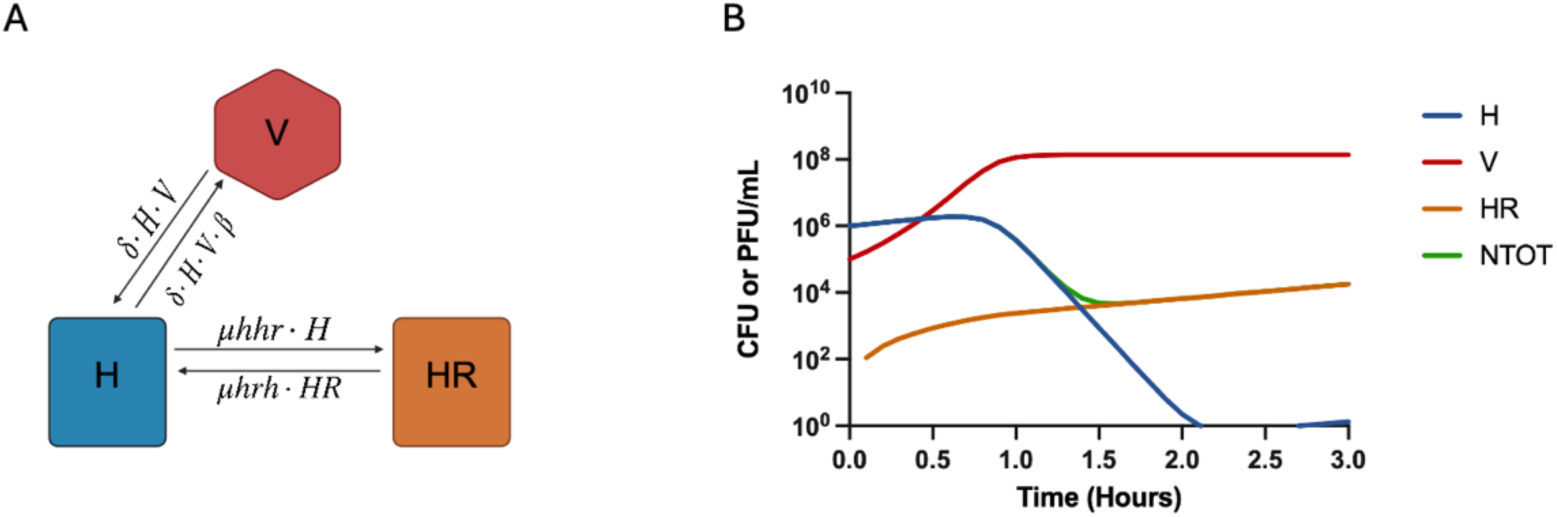
Diagram and Predictions of the model of heteroresistance. **(A)** Diagram of the heteroresistance model. The variables H, V, and HR are, respectively, sensitive heteroresistant bacteria, viral particles, and heteroresistant bacteria in cells or phage per mL. **(B)** Computer simulations of the proposed model of envelope resistance. H (sensitive heteroresistant bacteria, blue), V (viral particles, red), HR (heteroresistant bacteria, orange), NTOT (total bacteria, green).

### The heteroresistant population is unstable

To test whether reversion to sensitivity would occur in *E. coli* DS566-11 and *E. coli* DS566-15, we performed serial transfers without phage over 20 days^22^. Each transfer was spot tested with both phages, HP3 and ES19, to determine resistance (R) or susceptibility (S) as shown in Table 2 and Supplemental Table 6. After 14 days, *E. coli* DS566-15 reverted to susceptibility; however, *E. coli* DS566-11 did not revert within the timespan. As *E. coli* DS566-15 has a higher fitness cost than *E. coli* DS566-11 (Figure 2 and Supplemental Figure 4), consistent with stronger selection to shed a costly resistance mutation.

**Table 2.**
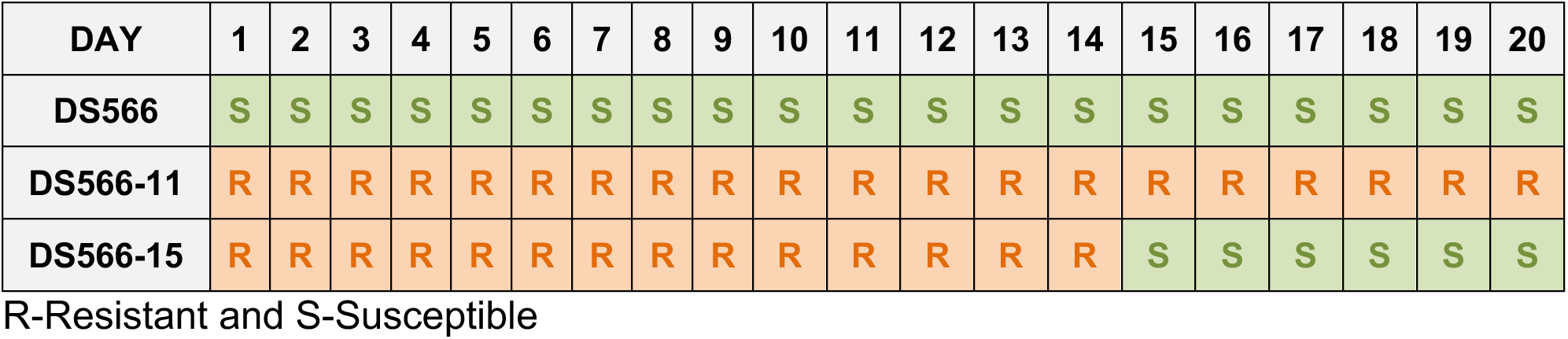
Reversion to susceptibility in urine.

### Whole-genome sequencing reveals convergent LPS-pathway mutations driving resistance

To elucidate the genetic architecture governing phage resistance and subsequent phenotypic reversion, we performed whole-genome sequencing (WGS) across a strategic panel of isolates. This cohort included the parental *E. coli* strain (DS566) alongside five derived variants representing distinct evolutionary trajectories: two time-kill-selected resistant mutants with contrasting fitness profiles (DS566-11, low cost; DS566-15, high cost), a phenotypically reverted derivative of the high-cost mutant (DS566-15R), and two independent resistant isolates recovered from fluctuation tests in LB broth (DS_HPLB) or human urine (DS_HPUrine). By comparative analysis of these stable, unstable, and independently evolved phenotypes, we sought to determine whether phage resistance and subsequent reversion are driven by parallel genetic mutations (Supplemental Text).

Initial pangenome comparisons confirmed that broad changes in gene content, such as gene gain or loss, do not explain the observed phenotypes.

Instead, high-resolution variant calling identified fixed, convergent coding lesions within the lipopolysaccharide (LPS) biosynthesis pathway unique to each resistant isolate (Supplemental Table 2). First, DS566-11 (stable, low-cost) carries a nonsense mutation in *rfaH*, predicted to disrupt the expression of the O-antigen and outer-core operons while leaving the inner core intact. Second, DS566-15 (unstable, high-cost) features a missense substitution in *hldE*, a key enzyme in ADP-heptose biosynthesis. Disruption of this locus is known to produce a severe, “deep-rough” LPS core deficiency, explaining its steep fitness penalty. Third, DS_HPLB and DS_HPUrine converged on the same pathway via a frameshifting deletion in *rfbC* (O-antigen biosynthesis) and a nonsense mutation in *rfaD* (a core-heptose epimerase), respectively.

No high-confidence, fixed mutations were detected in other standard outer-membrane protein receptor systems (e.g., *ompA*, *ompC*, *ompF*, *fhuA*, *lamB*) across any of the isolates.

Crucially, the phenotypic instability observed in the DS566-15 lineage was directly linked to the restoration of the wild-type allele. The susceptible revertant (DS566-15R) selectively lost the *hldE* mutation via back-mutation to the wild-type sequence, entirely relieving its associated fitness cost and restoring full phage susceptibility. This genomic divergence closely tracks with our phenotypic stability data; the low-cost mutant (DS566-11) did not experience strong selection for reversion and remained genetically and phenotypically stable throughout a 20-day serial passage (Table 2). Taken together, this striking pathway-level convergence across independent evolutionary replicates demonstrates that susceptibility to phages HP3 and ES19 is strictly dependent on intact, accessible host surface lipopolysaccharides.

## Discussion

Due to its specificity for targeting bacteria and its ability to replicate at the infection site, phage therapy is a promising therapeutic strategy for infectious diseases. Phage therapy is especially well-suited to bacterial infections such as UTIs, which present high loads of actively growing bacteria. However, as with antibiotics, resistance also evolves in these populations^23,24^. In this study, we present a particularly challenging form of resistance to phage: phage heteroresistance.

Heteroresistance is described as the phenomenon where subpopulations of seemingly isogenic bacteria exhibit a range of susceptibilities to antibiotics^20,25^. Antibiotic heteroresistance is traditionally tested for using a PAP test, by exposing different bacterial populations to increasing antibiotic concentrations to detect minority subpopulations^26^. Thus far, phage heteroresistance has not been described except for a recent clinical study conducted on a patient presenting with a recurrent case of *Bordetella bronchialis* infection; this patient received phage therapy and appeared to overcome the infection. However, by day eight, bacterial resistance rapidly took over, resulting in treatment failure on day nine^18^. Our results demonstrate that the heteroresistance we describe is phage dependent. We further find that this result is independent of growth medium, since it was consistent in both LB and urine.

The genetic basis of this heteroresistance helps explain why it arises so readily. Every resistant isolate we sequenced carried a loss-of-function lesion in the lipopolysaccharide (LPS) biosynthesis pathway, yet each arrived there through a different gene. Because both HP3 and ES19 depend on an intact LPS to adsorb, resistance can be achieved by disrupting any of the many genes required to build that surface, a large mutational target expected to generate resistant cells at high frequency, consistent with the elevated, phage-specific mutation rates and the sizable pre-existing resistant subpopulation we observed. The same architecture accounts for the instability of resistance: the depth of the LPS truncation tracks with its fitness cost, so that a severe, deep-rough lesion is both costly and readily reversed by back-mutation, whereas a milder lesion persists. Viewed this way, heteroresistance to these phages is an emergent property of targeting a receptor whose synthesis can be disrupted in many ways at different costs.

This phenomenon cannot be detected by the common susceptibility methods used, such as spot tests, plaque assays, and time kills^27–29^. For instance, a spot test can be misleading because it only shows whether a strain is susceptible or resistant; it does not account for heteroresistant subpopulations. When a spot test was performed on a heteroresistant strain such as *E. coli* DS566, it showed complete sensitivity to the phage. By contrast, when *E. coli* DS566 was co-incubated with the phage in a time kill assay, resistance appeared to evolve following a rapid decline of the total bacterial population. However, this result alone was not enough to separate it from classical resistance until mutation rates and mathematical models were considered. The limitations of the current methods used for phage susceptibility testing highlight the need for more accurate detection of resistant subpopulations.

As discussed earlier, the best method to determine heteroresistance is a PAP test, which has not yet been used for phage. Here we present a PAP test modified for the detection of phage heteroresistance analogous to the dilute-and-delay assay for antibiotic heteroresistance^30^. Unlike the current phage susceptibility methods, such as the spot test and time kill, this test directly reveals and quantifies minority resistant subpopulations. This is a key factor in determining whether a bacterial strain is heteroresistant.

The inability to properly detect phage heteroresistance can lead to treatment failure. In the case of UTIs, this can contribute to the high rates of recurrent UTIs^31^. Such a scenario was seen in a study that utilized a phage cocktail consisting of HP3 and ES19 to treat an *E. coli* UTI; the treatment appeared successful initially but the infection later recurred^14^. On a broader scale, studies have shown that treatment failure due to heteroresistance can contribute to the transmission of an infection^32,33^.

Our study has notable limitations. We examined a single clinical UPEC strain against two phages, so the generality of these findings to other host-phage pairs remains to be established, though the convergence on LPS across independently derived isolates suggests the underlying principle is not strain-specific. The experiments were performed in vitro; while pooled human urine provides a clinically relevant medium, it cannot capture the immune, anatomical, and pharmacokinetic factors that shape phage therapy in patients. The stability and reversion analyses rested on a small number of isolates, and we inferred the genetic basis of resistance from whole-genome sequencing and phenotype rather than from targeted genetic reconstruction. Finally, our phage PAP test is a proof of concept; standardization and prospective clinical validation will be needed before it can guide phage selection in practice.

Effective phage treatment can become more difficult to achieve when phage heteroresistance is present. In phage therapy, this is an important factor to consider when choosing a phage for treatment. Further research on this subject is essential to fully understand its mechanism and to develop effective strategies for its management in clinical care. The first step is proper *in vitro* testing that encompasses susceptibility, resistance, and heteroresistance. Our PAP test reveals the spectrum of phage susceptibility within a bacterial strain. To our knowledge, our study is the first formal demonstration of phage heteroresistance for UTIs in vitro. Taken together, our results offer one plausible explanation for the higher than anticipated rate of treatment failure for phage therapy: heteroresistance.

## MATERIALS AND METHODS

### Growth media

All experiments were conducted in pooled human urine and Luria-Bertani broth. Urine was collected from six participants that have not had antibiotics in the last six months, then sterilized using vacuum filtration.

### Growth conditions

All experiments were conducted at 37°C with shaking.

### Bacterial strain

All experiments were performed using the clinical UPEC strain *E. coli* DS566, which was isolated from patients with spinal cord injury.

### Bacteriophages

Phages HP3 (KY608967) and ES19 (MN508616) were isolated from environmental sources and wastewater, respectively.

### Spot test

*E. coli* DS566 was incubated overnight in either LB or urine media. One hundred μL of *E. coli* DS566 was added to 4 mL of soft agar, vortexed, and poured onto a phage plate. After five minutes, 10 μL of phage lysate is spotted onto the plate and then placed in the incubator at 37°C overnight.

### Time kill experiments

*E. coli* DS566 was incubated overnight in either LB or urine media. After overnight incubation, *E. coli* DS566 was diluted to 10^7^ CFU per mL and mixed with HP3 or ES19 diluted to 10^6^ PFU per mL in fresh LB or urine media. Viable bacterial cells and phage densities were estimated at 0, 10, 20, 30, 40, 50, 60, 90, 120, and 180 minutes. Experimental cultures were incubated overnight to estimate densities after 24 hours.

### Fluctuation tests

For the phage fluctuation test experiments, independent overnights of *E. coli* DS566 or *E. coli* A16 were exposed to either HP3 or ES19 for one hour before being plated on phage plates. For the antibiotic fluctuation tests, independent overnights of *E. coli* DS566 were plated on LB agar plates containing 5x MIC of streptomycin. Overnights of *E. coli* A16 were plated on 5x MIC of streptomycin. Experiments were performed with 20 biological replicates, and the mutation rates were calculated with BZrates.com.

### Growth curve estimates and suppression index

Exponential growth rates were estimated from changes in optical density (OD600) in a Bioscreen C. For this, 24-hour overnight cultures were diluted in either LB or urine media to an initial density of approximately 10^5^ cells per mL. To calculate the suppression index, *E. coli* DS566 was challenged with either HP3 or ES19 at an initial MOI of 10 and then serially diluted ten-fold four times. Five technical replicates were performed for each condition in a 100-well plate. The plates were incubated at 37°C and shaken before and after each measurement. Estimates of the OD600 were made every 5 minutes for 24 hours. Normalization was performed and means and standard deviations of the maximum growth rate, lag time, and maximum OD were found using an R Bioscreen C analysis tool accessible at https://josheclf.shinyapps.io/bioscreen_app.

### Serial transfers

Overnight cultures of *E. coli* DS566, *E. coli* DS566-11 and *E. coli* DS566-15 were diluted in either LB or urine media to approximately 10^5^ cells per mL in fresh media and incubated overnight. This transfer was performed every day for a total of twenty days. Each day overnight cultures were spot tested with HP3 and ES19 on phage plates.

### Population analysis profile tests

On 96-well plates 10^5^ PFU per mL of either HP3 and ES19 and 10^7^ bacterial cells per mL were added to each well to a total of 250 µL of LB or urine media. The cultures were then diluted ten-fold until 10 cells per mL were present in each well. The 96-well plates were incubated overnight. The turbidity of the wells was determined using spectrometry at 630 nm.

### Statistical analysis

Statistical significance analysis was carried out by paired t-tests and area under the curve using GraphPad Prism (version 10.2.0).

### Numerical Solutions (Simulations)

For our numerical analysis we used Berkeley Madonna with the parameters presented in Supplemental Table 4 and 5. Copies of the Berkeley Madonna programs used for these simulations are available at www.eclf.net.

### Sequencing

#### Genome sequencing, assembly, and annotation

Genomic DNA was extracted from single colonies and sequenced on an Illumina NovaSeq X Plus platform (SeqCenter). Reads were adapter-trimmed and quality-filtered, and draft genomes were assembled with Unicycler v0.5.0. Assembly quality was assessed with QUAST v5.2.0; genomes had a median (range) of 216 (112–260) contigs and a median (range) N50 of 299,517 (233,309-362,063) bp. Genes were predicted and annotated with Bakta v1.8.1 (database v5.0, full).

#### Pangenome analysis

Annotated assemblies were compared with Panaroo v1.7.0 to construct a pangenome and a gene presence/absence matrix. Custom Python scripts tested for gene clusters that distinguished the resistant isolates from the parent and revertant in either direction (uniquely present or uniquely absent). Where Panaroo reported altered local gene models at candidate loci, the underlying alignments were inspected to distinguish true gene loss from nucleotide-level disruption or contig-boundary fragmentation.

#### Read mapping and variant calling

The parental DS566 draft assembly was used as the reference for read-mapped variant calling. Paired-end Illumina reads from DS566 and each comparison isolate (DS566-11, DS566-15, DS566-15R, DS_HPLB, DS_HPUrine) were mapped to the reference with minimap2 v2.30 using the short-read preset (-ax sr). Alignments were coordinate-sorted and indexed with samtools v1.23.1, and per-sample mapping quality (percent reads mapped via samtools flagstat; mean depth via samtools depth -a) was recorded (**Supplemental data**).

Variants were called with bcftools v1.23.1. bcftools mpileup was run against the DS566 reference with per-sample depth and allelic-depth tags (-a FORMAT/DP,FORMAT/AD), followed by haploid variant calling (bcftools call -mv--ploidy 1). Calls were filtered to non-reference variants with QUAL ≥ 20 and sample depth ≥ 10 and normalized against the reference. Variants were exported with chromosome, position, reference and alternate alleles, QUAL, DP, and AD.

#### Variant annotation and focal validation

Variants were annotated against the DS566 Bakta GenBank file using a custom Python script (annotate_variants.py; Python v3.11.15, Biopython v1.86). For each variant the script identified the overlapping CDS; assigned gene, locus, and product; classified substitutions as synonymous, nonsynonymous, stop-gain, or stop-loss; and classified indels as in-frame or frameshift from the length change modulo three, with amino-acid consequences computed on the coding strand (including reverse-complement handling for negative-strand genes). Focal resistance mutations were validated directly from BAM pileups with samtools mpileup v1.23.1 (-aa -d 10000 -f DS566.fasta -r contig:position-position), and a custom parser (extract_bam_locus_support.py) counted exact alternate bases for SNVs and exact deletion events for the *rfbC* indel. The same procedure was used to confirm the absence of the *hldE* G216D allele in the revertant DS566-15R.

## Acknowledgments

We would like to thank the other members of the Levin Lab. Funds for this research were provided by a grant from the US National Institute of General Medical Sciences via R35GM136407 and the US National Institute of Allergy and Infectious Diseases via U19 AI 158080-02. UTI strains and phages were provided by Dr. Anthony Maresso. The content of this article is solely the responsibility of the authors and does not necessarily reflect the official views of the NIH. The funders had no role in the design or execution of this research nor the decision to publish the results.

## Author Contributions

Conceptualization: ASI, TGG, BAB, BRL

Methodology: TGG, BAB, BRL

Investigation: SSW, HES

Visualization: SSW, TGG

Funding Acquisition: BRL

Project Administration: BRL

Supervision: ASI, TGG, BAB, BRL

Writing– Initial Draft: SSW, ASI, TGG, BAB, MHW, BRL,

Writing– Review & Editing: SSW, ASI, TGG, BAB, MHW, BRL

## Competing Interest Statement

The authors have no conflicts to declare.

## Notes

https://www.eclf.net

### Competing Interest Statement

The authors have declared no competing interest.

