## Supplementary material for "Bacteriophage Heteroresistance as a Cause of Treatment Failure in Urinary Tract Infections": pdf

#### Model of phage and bacteria with envelope resistance

There is one population of lytic phage and two populations of bacteria with densities (particles and cells per ml) and designations *V the phage, N* phage-sensitive bacteria, and *R* phage-resistant bacteria. The growth of the bacteria is limited by the concentration of the single resource, r (µg per mL). The maximum rate of growth of the N and R bacteria are given by *rn* and *rr* (per cell per hour). The bacterial populations transition between each state with mutation rates µnr and µrn (per hour). The phage can adsorb to and replicate on N with a rate constant $\delta$ (per cell per hour), and a burst size $\beta$ (particles per infected bacterium). The rates of growth and phage infection are proportional to the concentration of the resource, $\psi(r) = \frac{r}{(r+k)}$ where k is the Monod constant and r the concentration of the resource.

With these definitions and assumptions, the rates of change in the concentration of the resource, density of the phage and the different bacterial populations are given by a set of differential equations,

$$\frac{dr}{dt} = -e \cdot \psi(r) \cdot (rn \cdot N + rr \cdot R) \qquad \text{Equation 1}$$

$$\frac{dN}{dt} = rn \cdot N \cdot \psi(r) - \delta \cdot V \cdot N \cdot \psi(r) - \mu nr \cdot N + \mu rn \cdot R \qquad \text{Equation 2}$$

$$\frac{dNR}{dt} = rr \cdot R \cdot \psi(r) - \mu rn \cdot R + \mu nr \cdot R \qquad \text{Equation 3}$$

$$\frac{dV}{dt} = \delta \cdot N \cdot V \cdot \beta \cdot \psi(r) \qquad \text{Equation 4}$$

The parameters used in the simulations are in Supplemental Table 4.

**Model of Phage Heteroresistance**

There is one population of lytic phage and two populations of bacteria with densities (particles and cells per ml) and designations *V the phage, H* the phage-sensitive population which generate the phage-resistant bacteria, and *HR* the phage-resistant bacterial subpopulation. The growth of the bacteria is limited by the concentration of the single resource, r (µg per mL). The maximum rate of growth of the H and HR bacteria are given by *rh* and *rhr* (per cell per hour). The bacterial populations transition between each state with mutation rates µnr and µrn (per hour). The phage can adsorb to and replicate on H with a rate constant $\delta$ (per cell per hour), and a burst size $\beta$ (particles per infected bacterium). The rates of growth and phage infection are proportional to the concentration of the resource, $\psi(r) = \frac{r}{(r+k)}$ where k is the Monod constant, the concentration of the resource, where the growth rate is half its maximum value.

With these definitions and assumptions, the rates of change in the concentration of the resource, density of the phage and the different bacterial populations are given by a set of differential equations,

$$\frac{dr}{dt} = -e \cdot \psi(r) \cdot (rh \cdot H + rhr \cdot HR)$$ Equation 5

$$\frac{dH}{dt} = rh \cdot H \cdot \psi(r) - \delta \cdot V \cdot H \cdot \psi(r) - \mu hhr \cdot H + \mu hrh \cdot HR$$ Equation 6

$$\frac{dHR}{dt} = rhr \cdot HR \cdot \psi(r) - \mu hrh \cdot HR + \mu hhr \cdot H$$ Equation 7

$$\frac{dV}{dt} = \delta \cdot H \cdot V \cdot \beta \cdot \psi(r)$$ Equation 8

The parameters used in the simulations are in Supplemental Table 5.

### Extended Results: Genomic Architecture Analysis

#### Resistance is not explained by gene gain or loss

A pangenome comparison showed that the parental, resistant, and reverted isolates share a highly conserved core genome. No gene cluster was uniquely shared by the parent and revertant while absent from all resistant isolates, and no gene cluster was uniquely shared by all resistant isolates while absent from both the parent and revertant. Candidate envelope genes were retained across the panel. Where Panaroo identified altered local gene models, these reflected nucleotide-level disruption consistent with draft assembly fragmentation rather than whole-gene loss. Broad differences in gene content therefore do not explain the resistance or reversion phenotypes. We therefore analyzed genome-wide read-mapped variants to identify nucleotide-level changes whose allele frequency, predicted effect, pathway, and genotype-phenotype pattern could explain resistance and reversion.

#### Resistant isolates have convergent LPS-pathway lesions

Haploid read-mapped variant calling against the DS566 reference identified fixed coding lesions in LPS biosynthesis genes in each resistant isolate (Supplemental Table 2). We separately inspected allele-depth support in BAM pileups at additional envelope-associated loci, which revealed recurrent intermediate-frequency or lineage-associated *tsx* and *xylH* variants. However, these did not show the same genotype-phenotype concordance as the fixed LPS-pathway lesions. We therefore prioritized the fixed LPS-pathway lesions as the primary resistance candidates. The stable, low-cost mutant DS566-11 carries a nonsense mutation in *rfaH* (W4*), predicted to disrupt RfaHdependent expression of the O-antigen and outer-core operons while leaving the inner core intact, which is consistent with its low fitness cost.

The high-cost mutant DS566-15 carries a missense substitution in *hldE* (G216D), the bifunctional kinase/adenylyltransferase that synthesizes the ADP-heptose donor for the LPS inner core[1, 2]. Loss of HldE function yields a heptose-deficient, 'deep-rough' LPS associated with outer-membrane destabilization and reduced fitness[1, 2]. The high fitness cost we observe for DS566-15 (Figure 2) is consistent with such a mechanism, although we did not directly profile its LPS. The two independently selected fluctuation-test isolates converged on the same pathway by different routes: DS_HPLB carries a frameshifting deletion in *rfbC* (O-antigen/rhamnose biosynthesis), and DS_HPUrine carries a nonsense mutation in *rfaD* (Q244*, core-heptose epimerase). No high-confidence, fixed mutations were detected in other characterized phage-receptor systems, including *ompA*, *ompC*, *ompF*, *fhuA*, *btuB*, *lamb*, in any isolate[3]. Other variants were detected outside these loci, including recurrent *tsx* changes and lineage-associated *xylH* variants, but these did not show the same concordance with resistance and reversion. For example, *tsx* L14R and *xylH* G128R were retained in the susceptible revertant. Together, despite divergence in predicted receptor-binding tail-fiber proteins, the convergence of independent resistant isolates on LPS biosynthesis genes supports a model in which HP3 and ES19 susceptibility depends on intact LPS or LPS-mediated surface access.

<u>Phenotypic susceptibility reversion in DS566-15 is linked to restoration of wild-type *hldE*</u>

The instability of the DS566-15 phenotype is strongly linked to loss of *hldE* G216D. Loss of *hldE* G216D in the susceptible revertant without compensatory second-site mutations, together with retention of lineage-associated *tsx* and *xylH* variants, strongly implicates *hldE* G216D as the reversible resistance-associated allele in this lineage. Restoration of susceptibility required loss of only a single-nucleotide difference at *hldE*. Given published estimates of spontaneous *E. coli* base-substitution rates on the order of $10^{-10}$ mutations per nucleotide per generation[(4)], restoration of the wild-type allele is mutationally plausible during high-density serial passage, although we did not directly measure the reversion rate. This is consistent with the observation that the low-cost mutant DS566-11 did not revert within 20 days, and links the genomic result to the phenotypic instability of this lineage.

<u>Genotype–phenotype concordance</u>

Supplemental Table 2 summarizes the concordance between each isolate's mutation, its predicted molecular and LPS-level consequence, its measured fitness cost and phage phenotype, and whether the resistance allele is stable or unstable across 20-day serial passage. The O-antigen-associated candidates (*rfaH* in DS566-11 and *rfbC* in DS_HPLB) are predicted to disrupt distal LPS structures, whereas the core-biosynthesis candidates (*hldE* in DS566-15 and *rfaD* in DS_HPUrine) are predicted to produce rough or deep-rough LPS phenotypes. The reverting lineage (DS566-15) carries a point substitution in core biosynthesis whose back-mutation restores both LPS and susceptibility.

<u>Limitations</u>

There are three important limitations of this genomic analysis. First, variant calling used a fragmented draft assembly (median 216 contigs) as the reference, so variants near contig breaks, structural variants, and mobile-element rearrangements may be missed. Second, pangenome presence/absence on highly fragmented assemblies can produce spurious gene-model fragmentation at contig ends, so we relied on read-level evidence for all causal claims. Third, short-read data cannot reliably resolve homopolymeric-tract length variation or invertible elements; we found no evidence of phase variation at the resistance-associated loci, but phase variation elsewhere in the genome cannot be excluded without long-read sequencing.

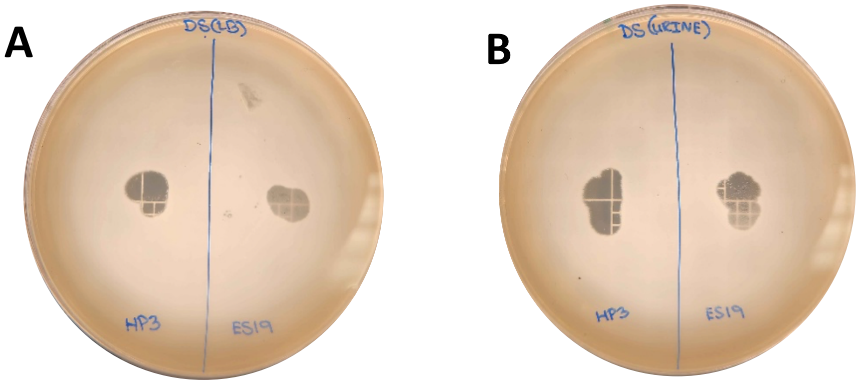


**Supplemental Figure 1. Spot test.** Spot tests with the phage HP3 (left side of plate) and the phage ES19 (right side of plate) on an *E. coli* DS566 lawn in LB **(A)** and urine media **(B)**

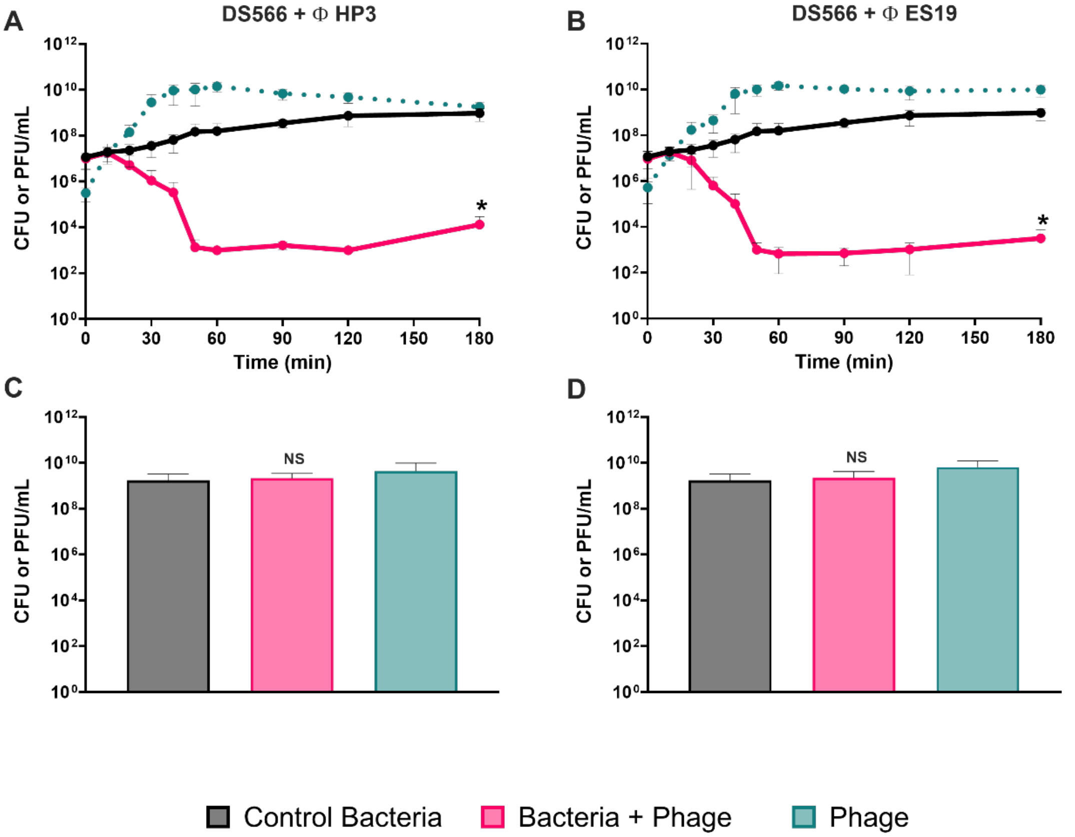


**Supplemental Figure 2.** ***E. coli* DS566 time kill in LB.** Shown at the top are three-hour time kills with *E. coli* DS566 and HP3 **(A)** or ES19 **(B)**. Shown below are the 24-hour densities from the time kill experiments **(C, D)**. In black is the bacterial density of a phage-free control, pink is the bacterial density of a culture with the phage, and blue is the phage density in the culture with the phage. Shown are means and standard deviations of three biological replicates. *$p < 0.1$, ns = not significant.

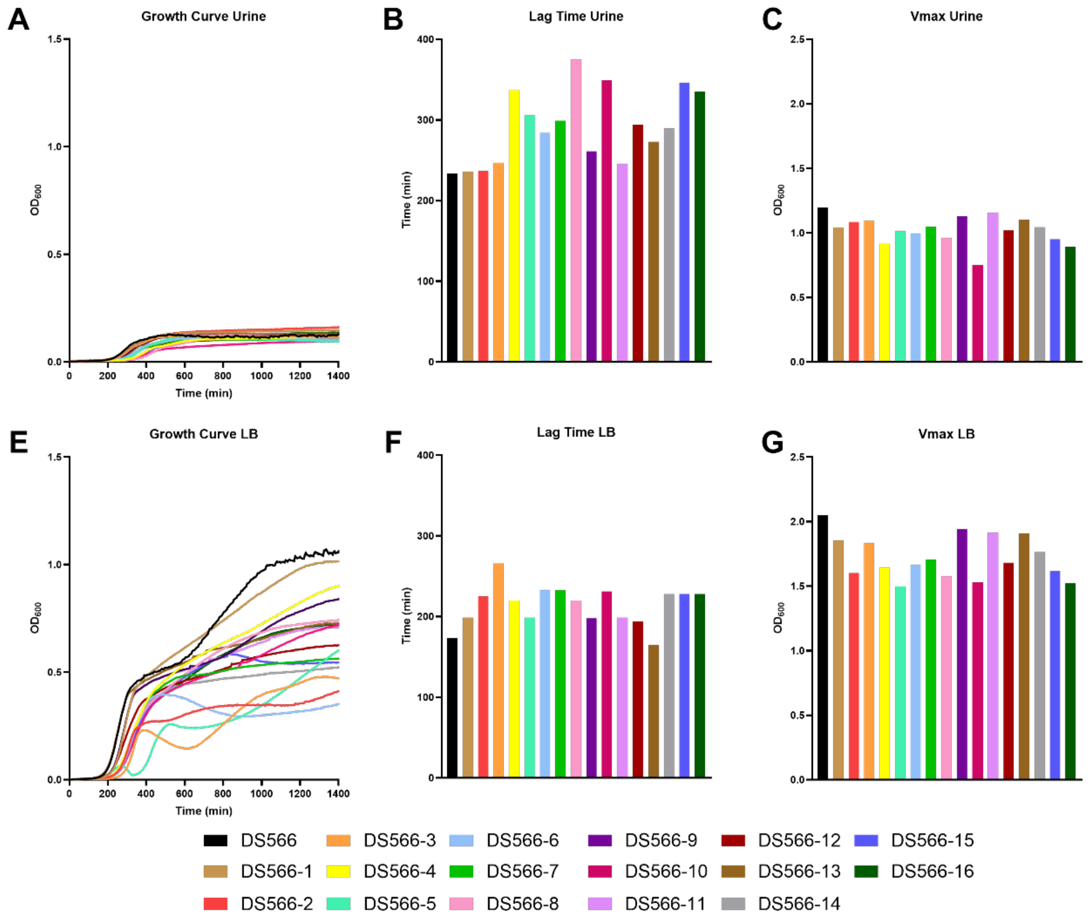


**Supplemental Figure 3. Growth dynamics of the *E. coli* DS566 resistant mutants.** Shown at the top are the dynamics in urine. Shown at the bottom are the dynamics in LB media. **(A,E)** mutant growth curves. **(B,F)** lag times of each mutant. **(C,G)** Vmax of each mutant.

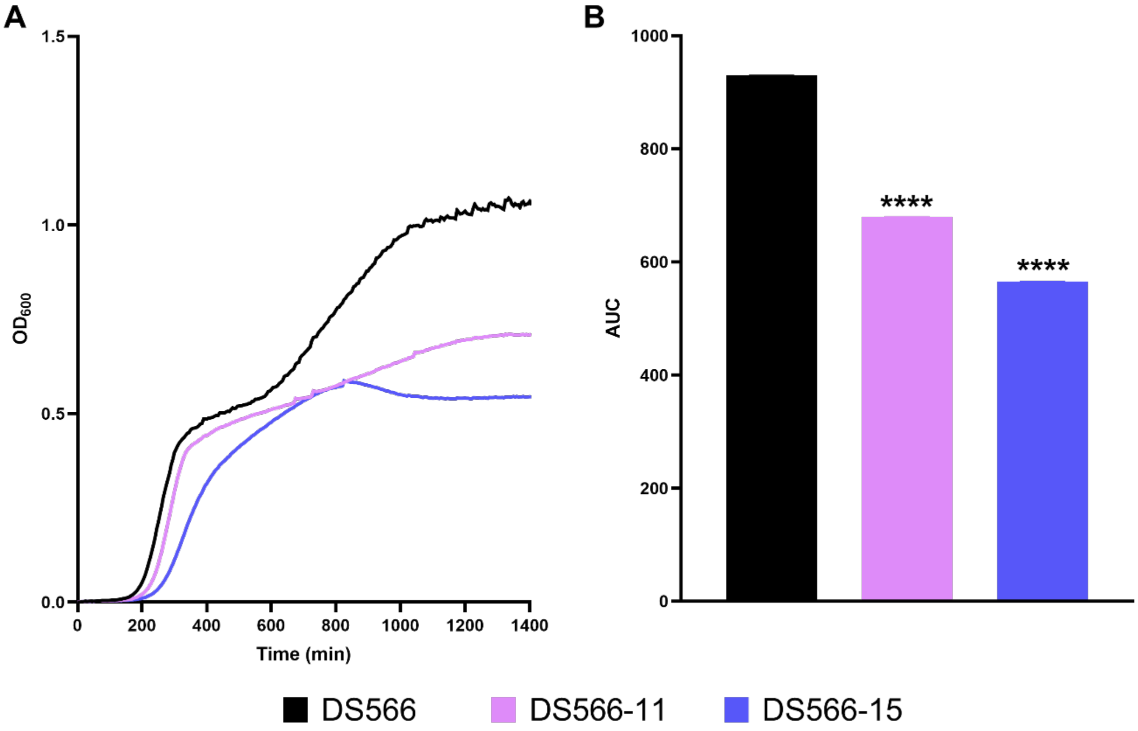


**Supplemental Figure 4. Fitness cost of phage-resistant mutants in LB.** Shown in black is the ancestral *E. coli* DS566, in pink a low-fitness-cost mutant, and in indigo a high-fitness-cost mutant. Bar graphs represent the means and standard errors of five technical replicates. **(A)** OD growth curves. **(B)** Area under the curve of the OD growth curves. ****$p$ < 0.0001, ns = not significant

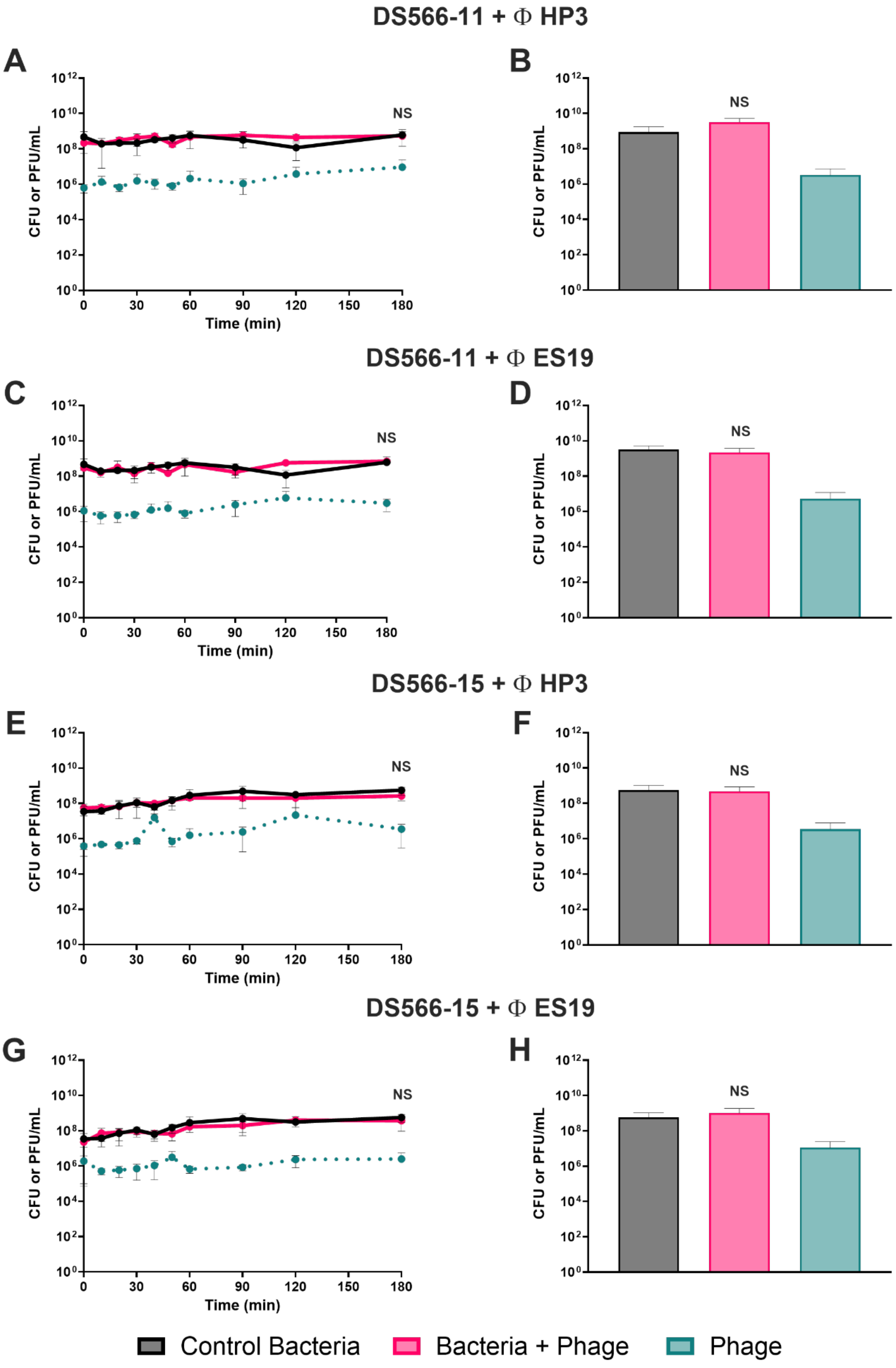
DS566-11 + Φ HP3
A
B
DS566-11 + Φ ES19
C
D
DS566-15 + Φ HP3
E
F
DS566-15 + Φ ES19
G
H
CFU or PFU/mL
Time (min)
NS
Control Bacteria
Bacteria + Phage
Phage

**Supplemental Figure 5. Time kills of phage-resistant mutants with either phage**

Time kills with the phage-resistant mutants and either phage. **(A)** Time kill with *E. coli* DS566-11 with the phage HP3. **(B)** 24-hour densities of *E. coli* DS566-11 with the phage HP3. **(C)** Time kill with *E. coli* DS566-11 with the phage ES19. **(D)** 24-hour densities of *E. coli* DS566-11 with the phage ES19. **(E)** Time kill with *E. coli* DS566-15 with the phage HP3. **(F)** 24-hour densities of *E. coli* DS566-15 with the phage HP3. **(G)** Time kill with *E. coli* DS566-15 with the phage ES19. **(H)** 24-hour densities of *E. coli* DS566-15 with the phage ES19. ns = not significant. In black is the bacterial density of a phage-free control, pink is the bacterial density of a culture with the phage, and blue is the phage density in the culture with the phage. Shown are means and standard deviations of three biological replicates.

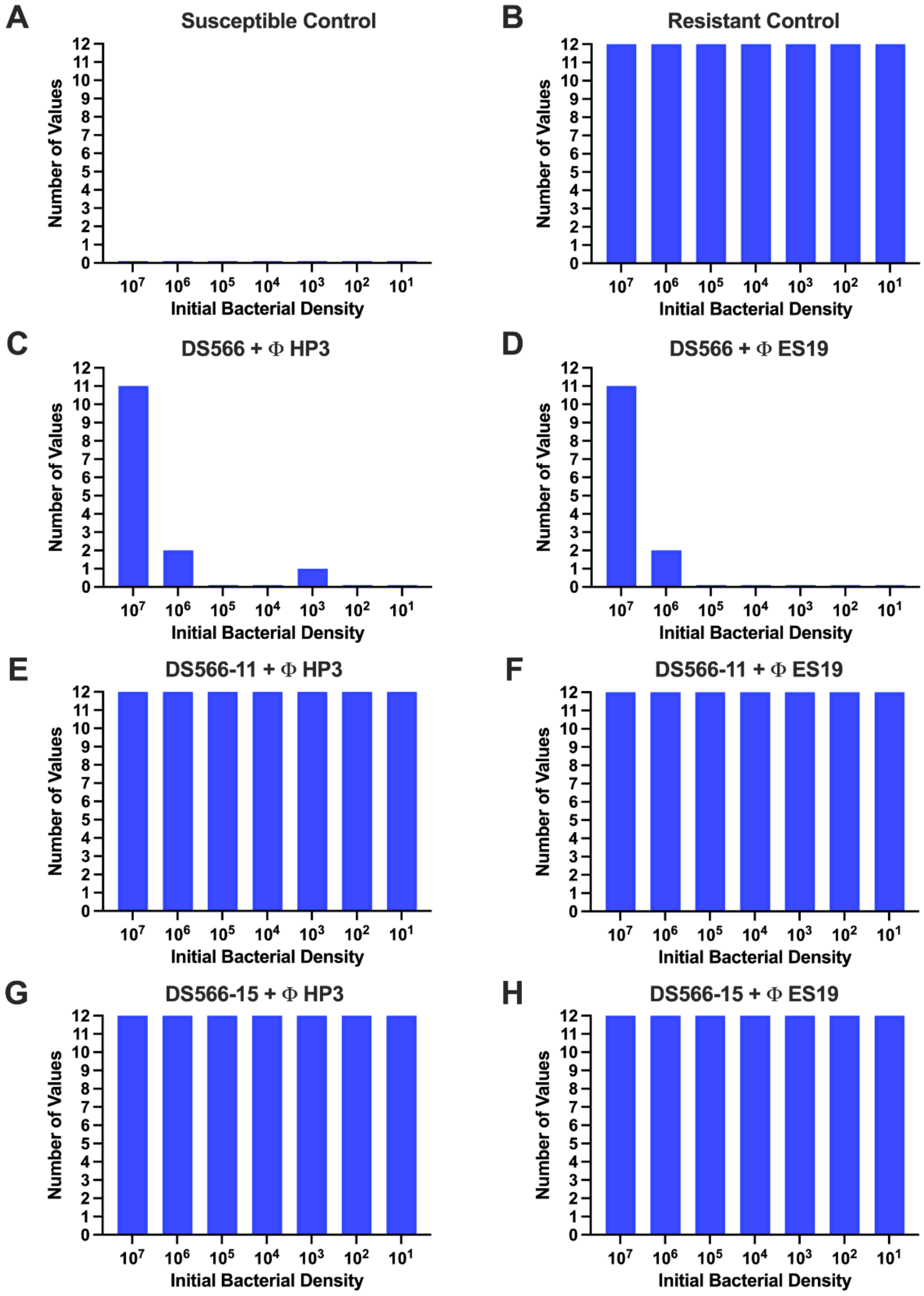
A
Susceptible Control
B
Resistant Control
C
DS566 + Φ HP3
D
DS566 + Φ ES19
E
DS566-11 + Φ HP3
F
DS566-11 + Φ ES19
G
DS566-15 + Φ HP3
H
DS566-15 + Φ ES19
Number of Values
Initial Bacterial Density
10^7 10^6 10^5 10^4 10^3 10^2 10^1

**Supplemental Figure 6. Phage PAP test in LB. (A)** *E. coli* C with the phage T6 (susceptible control). **(B)** *E. coli* DS566 with the phage T3 (resistant control). **(C)** *E. coli* DS566 with the phage HP3. **(D)** *E. coli* DS566 with the phage ES19. **(E)** *E. coli* DS566-11 with the phage HP3. **(F)** *E. coli* DS566-11 with the phage ES19. **(G)** *E. coli* DS566-15 with the phage HP3. **(H)** *E. coli* DS566-15 with the phage ES19.

**Supplemental Table 1. Spot tests of *E. coli* DS566 isolates at different timepoints in the time kill.**

| DS566 Mutants | Phage Isolated From | Media Isolated From | HP3 & ES19 Susceptibility | Timepoint Isolated in Time Kill (Hours) |
|---|---|---|---|---|
| **DS566-1** | HP3 | Urine | NO | 2 |
| **DS566-2** | HP3 | Urine | NO | 1.5 |
| **DS566-3** | HP3 | Urine | NO | 2 |
| **DS566-4** | HP3 | LB | NO | 2 |
| **DS566-5** | ES19 | Urine | NO | 1 |
| **DS566-6** | ES19 | Urine | NO | 1.5 |
| **DS566-7** | ES19 | Urine | NO | 1.75 |
| **DS566-8** | ES19 | LB | NO | 2 |
| **DS566-9** | HP3 | Urine | NO | 1 |
| **DS566-10** | HP3 | Urine | NO | 1.5 |
| **DS566-11** | HP3 | Urine | NO | 2 |
| **DS566-12** | HP3 | LB | NO | 2 |
| **DS566-13** | ES19 | Urine | NO | 1 |
| **DS566-14** | ES19 | Urine | NO | 1.5 |
| **DS566-15** | ES19 | Urine | NO | 1.75 |
| **DS566-16** | ES19 | LB | NO | 2 |

**Supplemental Table 2. Genotype–phenotype concordance for candidate resistance mutations in DS566-derived isolates.**

| Isolate (phenotype) | Gene/ mutation | Predicted effect | LPS pathway step | Predicted LPS phenotype/ fitness cost | Phage phenotype & allele stability |
|---|---|---|---|---|---|
| DS566-11 *(resistant)* | *rfaH* W4* | Stop-gain (nonsense at codon 4) | O-antigen + outer-core operon expression (RfaH antiterminator) | O-antigen loss, core intact / low cost | Resistant; stable (no reversion in 20 days) |
| DS566-15 *(resistant)* | *hldE* G216D | Nonsynonymous in HldE | ADP-heptose biosynthesis (LPS inner core) | Deep-rough core / high cost | Resistant; unstable (reverts by day 14) |
| DS566-15R *(revertant)* | *hldE* wild-type (G216 restored) | Back-mutation to WT allele | ADP-heptose biosynthesis restored | LPS restored / cost relieved | Susceptible; resistance allele lost; lineage markers retained |
| DS_HPLB *(HP3, LB)* | *rfbC* ~ 10-bp del | Frameshift → early truncation | dTDP-L-rhamnose / O-antigen biosynthesis | O-antigen loss, core intact / low–moderate cost | Resistant; stable (independent selection) |
| DS_HPUrine *(HP3, urine)* | *rfaD* Q244* | Stop-gain (truncates C-terminal ~22%) | ADP-heptose epimerase (LPS core) | Rough/deep-rough core / moderate–high cost | Resistant; stable (independent selection) |

**Supplemental Table 3. Summary of statistical significance for Luria-Delbrück tests**

| Mutation Rate | Vs | Mutation Rate | P < 0.05 |
|---|---|---|---|
| DS566 + HP3 | Vs | A16 + HP3 | NO |
| DS566 + ES19 | Vs | A16 + ES19 | NO |
| DS566 + HP3 | Vs | DS566 + Streptomycin | YES |
| DS566 + ES19 | Vs | DS566 + Streptomycin | YES |
| DS566 + HP3 | Vs | DS566 + ES19 | NO |
| A16 + HP3 | Vs | A16 + ES19 | NO |
| A16 + HP3 | Vs | A16 + Streptomycin | YES |
| A16 + ES19 | Vs | A16 + Streptomycin | YES |
| DS566 + Streptomycin | Vs | A16 + Streptomycin | NO |

**Supplemental Table 4. Parameters and the values used in the simulation of envelope resistance model**

| Parameter | Definition | Value | Units | Source |
|---|---|---|---|---|
| r | Maximum resource concentration | 1000 | µg/mL | This report |
| rn | Maximum growth rate N | 1 | per cell per hour | This report |
| rr | Maximum growth rate R | 1 | per cell per hour | This report |
| $\delta$ | Phage adsorption rate | $10^{-7}$ | per hour per mL | (5) |
| $\beta$ | Phage burst size | 50 | particles per cell | (5) |
| $\mu$nr | Mutation rate to resistant | $9.44 \times 10^{-7}$ | per hour | This report |
| $\mu$rn | Mutation rate to sensitive | $9.44 \times 10^{-7}$ | per hour | This report |
| k | Monod constant | 1.0 | µg | (6) |
| e | Resource conversion efficiency | $5 \cdot 10^{-7}$ | µg/cell | (7) |

**Supplemental Table 5. Parameters and the values used in the simulation of the joint action of heteroresistance model**

| Parameter | Definition | Value | Units | Source |
|---|---|---|---|---|
| r | Maximum resource concentration | 1000 | µg/mL | This report |
| rh | Maximum growth rate H | 1.2 | per cell per hour | This report |
| rhr | Maximum growth rate HR | 1 | per cell per hour | This report |
| $\delta$ | Phage adsorption rate | $10^{-7}$ | per hour per mL | (5) |
| $\beta$ | Phage burst size | 50 | particles per cell | (5) |
| $\mu$hhr | Transition rate to heteroresistant | 0.001 | per hour | This report |
| $\mu$hrh | Transition rate to sensitive | 0.001 | per hour | This report |
| k | Monod constant | 1.0 | µg | (6) |
| e | Resource conversion efficiency | $5 \cdot 10^{-7}$ | µg/cell | (7) |

**Supplemental Table 6. Reversion to sensitivity in LB**

| DAYS | 1 | 2 | 3 | 4 | 5 | 6 | 7 | 8 | 9 | 10 | 11 | 12 | 13 | 14 | 15 | 16 | 17 | 18 | 19 | 20 |
|---|---|---|---|---|---|---|---|---|---|---|---|---|---|---|---|---|---|---|---|---|
| DS566 | S | S | S | S | S | S | S | S | S | S | S | S | S | S | S | S | S | S | S | S |
| DS566-11 | R | R | R | R | R | S | R | S | R | R | R | R | R | R | S | R | R | R | R | R |
| DS566-15 | R | R | R | R | R | R | S | S | S | S | S | S | S | S | S | S | S | S | S | S |

R-Resistant and S-Susceptible
